# Impact of Low-Dose Narrowband UV Radiation on Skin Tissue and Mast Cells in a Canine Model

**DOI:** 10.1101/2025.01.01.631029

**Authors:** Yusuke Maeda, Shota Kubota, Shinpei Kawarai, Mai Kamimura, Hiroki Okuyama, Moeko Ono, Moe Nishida, Shigeharu Moriya, Yushi Onoda, Tomohiro Tsurumoto, Koichi Nagata, Shin-ichi Ansai, Yasuo Fujikawa, Atsushi Tsukamoto

**Author notes:** Corresponding authors E-mail (AT) E-mail (SK).

## Abstract

Narrowband ultraviolet radiation (NB-UVR) therapy is widely utilized in dermatology worldwide. While the immunosuppressive effects of ultraviolet radiation are well-documented, its non-inflammatory effects remain poorly understood. This study evaluated the effects of sub-minimal erythema dose (MED) narrowband ultraviolet radiation using a canine model. The effects of narrowband-ultraviolet B (310 nm) and A2 wavelengths (320 and 330 nm) were compared. Initially, we demonstrated the therapeutic potential of NB-UVR in dogs with spontaneous atopic dermatitis (n = 6, p < 0.05). Following 300 mJ/cm² sub-MED ultraviolet radiation exposure, intradermal skin testing, skin mast cell counts, and RNA sequencing analysis were conducted. Additionally, cell viability (WST assay) and apoptosis (DAPI staining) were assessed in vitro using a canine mast cell line. Ultraviolet radiation exposure significantly reduced intradermal skin test scores across all wavelengths and decreased skin mast cell counts at 310 nm (n = 3, p < 0.05). In vitro, cell viability was significantly reduced at 310 and 320 nm (p < 0.05), and 310 nm irradiation specifically increased apoptotic cell numbers (p < 0.05). RNA sequencing identified 71 differentially expressed genes at 320 nm, with enrichment analysis highlighting muscle cell components and functions, potentially associated with arrector pili muscle activity. These findings suggest that 310 nm irradiation induces apoptosis, while 320 nm elicits non-inflammatory modulatory effects, highlighting distinct wavelength-specific therapeutic mechanisms of narrowband ultraviolet radiation.

## Introduction

Ultraviolet radiation (UVR) from sunlight is essential for maintaining homeostasis in living organisms, playing key roles in cell growth, differentiation, melanogenesis, and vitamin D synthesis [1, 2]. However, excessive UVR exposure is associated with DNA damage, leading to skin carcinogenesis and premature aging. In recent decades, various immunomodulatory UV wavelengths have been developed for therapeutic applications, including UVA2 (315–340 nm), UVA1 (340–380 nm), broadband UVB (280–315 nm), and narrowband (NB) UVB (308–313 nm). These treatments have shown efficacy in managing conditions such as psoriasis, atopic dermatitis (AD), urticaria, mastocytosis, mycosis fungoides, and vitiligo [1–3]. Efforts to enhance safety and efficacy have focused on identifying precise UV wavelengths and developing narrower-bandwidth illuminators [2, 3]. Recent advancements, such as excimer lamps (308 nm) and light-emitting diode (LED) technologies, have refined the application of targeted phototherapy [3]. However, the specific therapeutic effects of narrower wavelengths, such as those separated by 10-nm intervals, remain insufficiently understood.

The mechanisms underlying the effects of therapeutic UVR doses have been extensively studied, particularly the keratinocyte damage pathway responsible for immunosuppression. UVR is absorbed by chromophores such as urocanic acid, nucleotide bases, and 7-dehydrocholesterol, which subsequently activate keratinocytes and immune cells, including Langerhans cells and mast cells (MCs) [1, 3]. These activated cells produce inflammatory (TNF), Th2 (IL-4), and anti-inflammatory (IL-10) cytokines, reducing antigen-presenting cell function, altering regulatory T cell populations, and suppressing systemic contact hypersensitivity (CHS) [1–3]. Additionally, UV-damaged necrotic keratinocytes release damage-associated molecular patterns (DAMPs). Components of these DAMPs, such as DNA and dsRNA, stimulate innate immune responses in neighboring keratinocytes, creating an environment conducive to repair and resistance against infections [1]. Since the early 2000s, studies have revealed the effects of UV-induced immunosuppression on MCs and their mediators, using models such as bone marrow-derived MC transplants into MC-deficient mice [4–7]. These UV-induced pathways are considered to orchestrate the overall responses. While these pathways are considered integral to the overall response, research has predominantly focused on UVR doses above the minimal erythema dose (MED), leaving the effects of sub-MED UVR largely unexplored [1, 8, 9]. Because MED UVR induces erythema, we hypothesized that sub-MED UVR would allow us to investigate its non-inflammatory effects.

Most prior studies of NB-UVR therapy have concentrated on therapeutic wavelengths for humans, often relying on mouse models. However, there is ongoing debate about whether therapeutic wavelengths optimized for humans are equally suitable for mice, given differences in skin structure. *Ex vivo* studies have also been reported [10], but these are potentially confounded by factors such as altered blood flow and surgical tissue damage. Dogs provide a valuable natural model for studying AD [11], offering several advantages over rodents. Canine MCs, which play a central role in canine AD, are biologically similar to their human counterparts in origin and in the expression of serine proteases such as tryptase and chymase [11, 12]. Furthermore, dogs develop AD spontaneously, presenting a more representative system for studying NB-UVB therapy [13, 14].

In this study, we examined the effects of sub-MED NB-UVR exposure on canine skin, including both healthy dogs and those with spontaneous AD. We compared the responses across different wavelengths, leveraging the thinner skin of dogs relative to humans to gain insight into the differential effects of UVR wavelengths.

## Material and Methods

### Animals

This study was approved by the Institutional Animal Care and Use Committee of Azabu University (approval number: 180924-5) and conducted in compliance with the ARRIVE guidelines 2.0 and the Guide for the Care and Use of Laboratory Animals by the National Institutes of Health. A total of 12 dogs were included in the study. Six client-owned dogs with spontaneous atopic dermatitis (AD) were enrolled in clinical trials of narrowband ultraviolet radiation (NB-UVR) therapy during Oct 1, 2018 to Jan 31, 2020 (S1 Table). Canine AD was diagnosed based on Favrot’s criteria [15]. Dogs with other causes of pruritus, such as primary bacterial or fungal infections, ectoparasitosis, flea bite hypersensitivity, or adverse food reactions, were excluded. All enrolled dogs presented with at least four skin lesions characterized by alopecia and dermatitis. Informed consent was obtained from all owners before the start of the clinical trials.

The remaining six dogs were healthy Beagles (2 males and 4 females; mean age: 5.8 years) obtained from Kitayama Labes (Nagano, Japan). These dogs were used to evaluate the effects of sub-MED repeated NB-UVR irradiation on immediate anaphylactoid reactions, histological changes, and gene expression in the skin. The Beagles were housed under suitable conditions, including safe, clean, and spacious environments with appropriate feeding and temperature controls, at the institute’s laboratory animal facility. Routine physical examination and blood tests were performed to maintain their health status.

### NB-UVR LED devices

The study utilized LED devices with peak wavelengths of 310 nm, 320 nm, and 330 nm for irradiation. All LED devices had a half-bandwidth of 10 nm (S1 Fig). The intensity on the irradiated surface was standardized at 6.5 mW/cm².

### Effects of NB-UVRs on dogs with spontaneous AD

The skin lesions of six client-owned dogs with spontaneous AD were treated with 310 nm or 320 nm NB-UVR irradiation. As a positive control, topical betamethasone valerate ointment (RINDERON-VG Ointment, Shionogi, Osaka, Japan) was applied, while untreated lesions served as the negative control (S2 Table). For NB-UVR treatments, an equal number of lesions were selected per dog, with one lesion designated for both positive and negative control treatments.

Dogs receiving systemic treatments (e.g., cyclosporine, antibiotics, antifungal drugs), undergoing shampoo or diet changes within two weeks before the trial, receiving vaccinations within two weeks, or with a history of malignant diseases were excluded. Treatments were administered weekly or biweekly, based on the owners’ schedules. The initial NB-UVR dose was set at 300 mJ/cm², determined to be below the MED range (400–1500 mJ/cm², n=6) based on preliminary experiments (S3 Table). The dose was increased by 100 mJ/cm² at each follow-up visit. Betamethasone ointment was applied at 0.05–0.1 g per lesion (approximately one fingertip unit).

Lesion severity was evaluated at each visit using the Canine Atopic Dermatitis Extent and Severity Index (CADESI-4) lesion grading atlas [16]. Four clinical signs (erythema, alopecia, excoriation, and lichenification) were scored on a six-point scale (0 = none, 1–2 = mild, 3 = moderate, 4–5 = severe), and the total lesional score was calculated by summing the scores for each sign (range: 0–20). For dogs with more than two NB-UVR-treated lesions, the average score was used. Percentage changes in lesional scores relative to baseline (week 0) were calculated. The scores for NB-UVR treatments (310 nm and 320 nm) were evaluated in a double-blinded manner.

### Intradermal skin test

The intradermal skin test (IDT) was conducted on Beagles under sedation with medetomidine (0.02 mg/kg, IM; Dorbene Vet, Kyoritsu Seiyaku, Tokyo, Japan) and midazolam (0.3 mg/kg, IM). Prior to the IDT, NB-UVR (310, 320, and 330 nm) was applied to hair-clipped skin at 300 mJ/cm² once daily for four consecutive days at the same sites, beginning at 13:00 each day. On the day following the final irradiation, the IDT was performed at 12 sites (three NB-UVR-irradiated sites and one unirradiated site) across the right lateral, back, and left lateral regions in triplicate.

Reagents (50 µL) were serially diluted 1:10 in saline and injected intracutaneously. Polyoxyethylene hydrogenated castor oil 60 (HCO-60; Kao Chemicals, Tokyo, Japan) was used as a mast cell (MC) degranulation indicator. Concentrations of 1 × 10⁻⁶ v/v (0.001%) and 1 × 10⁻⁵ v/v (0.01%) were selected based on the previously established threshold for MC degranulation in healthy dogs [17]. Histamine (1 × 10⁻⁵ w/v; Sigma Aldrich, St. Louis, MO, USA) served as the positive control, while saline was used as the negative control.

Photographs were taken of both pre- and post-IDT sites to assess changes in redness values (RV), and the maximum diameter (D) of wheal-and-flare formation was measured 15 minutes after injection. The RV was analyzed using computer software (GNU Image Manipulation Program, https://www.gimp.org/) by calculating mean red (R) and green (G) intensity (range: 0–256) using the color picker in RGB mode. RV was determined using the formula (R−G)/(R+G), and changes in RV were expressed as the ratio of post-IDT RV to pre-IDT RV. The IDT score was calculated as follows:

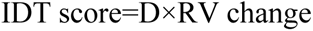

Measurements of D and RV were conducted in a blinded manner. Following the procedure, atipamezole (0.08 mg/kg; Atipame Injection, Kyoritsu Seiyaku) was administered intramuscularly to reverse sedation.

### NB-UVR irradiation and skin biopsy for histopathological analysis and RNA-sequencing

NB-UVR (310, 320, and 330 nm) was applied to the back region of Beagles at a sub-MED dose of 300 mJ/cm² once daily for four consecutive days, following the same protocol as described for the IDT. As an inflammatory control, a higher dose of NB-UVB (310 nm, 1500 mJ/cm²) was applied one day prior to the skin biopsy.

Skin biopsies were collected under sedation with medetomidine and midazolam. Tissue samples were obtained from irradiated sites using an 8-mm biopsy punch. Each sample was divided into two halves: one fixed in 9 mL of formalin, methanol, and picric acid-based fixative (Gfix; Genostaff, Tokyo, Japan) for histopathological analysis, and the other preserved in RNA later solution (Thermo Fisher Scientific, Tokyo, Japan) for RNA sequencing (RNA-seq). Fixed tissues were embedded in paraffin, sectioned at 5-μm thickness, and stained with hematoxylin and eosin (HE) or toluidine blue (TB, pH 4.1; Muto Pure Chemicals, Tokyo, Japan). These sections were analyzed for the presence of sunburn cells and MCs. RNA later-preserved tissues were sent to Takara Bio (Shiga, Japan) for RNA-seq analysis.

### Histopathological analysis

Epidermal sunburn cells were identified using HE-stained sections. MC counts were performed on TB-stained sections, focusing on a depth of 0.2 mm from the superficial dermis to the upper reticular dermis. The average number of MCs per section was calculated from four sections per sample. All histopathological evaluations were conducted in a blinded manner.

### Whole transcriptome RNA sequencing

RNA-seq analysis was performed by Takara Bio. Total RNA was extracted using the RNAiso Plus kit (Takara Bio) and further purified with the NucleoSpin RNA Clean-up XS kit (Takara Bio). RNA concentration and quality were evaluated using a NanoDrop spectrophotometer (Thermo Fisher Scientific) and the Agilent 2200 TapeStation (Agilent Technologies, Santa Clara, CA, USA). The average total RNA yield, 260/280 ratio, and 260/230 ratio (mean ± SD) were 14.1 ± 2.9 μg, 2.1 ± 0.0, and 2.0 ± 0.1, respectively.

Sequence libraries were prepared using the TruSeq Stranded mRNA Sample Prep Kit (Illumina, San Diego, CA, USA) and indexed with the IDT for Illumina TruSeq UD Indexes (Illumina) on an Agilent XT-Auto System (Agilent Technologies). For each sample, 2 μg of RNA was used for PolyA+ RNA isolation. Adaptor-ligated double-stranded DNA was amplified with 15 cycles of PCR. Sequencing was conducted on the NovaSeq 6000 system (Illumina) using the NovaSeq 6000 S4 Reagent Kit and NovaSeq Xp 4-Lane Kit for 150-bp paired-end reads. Data were processed with NovaSeq Control Software v1.4.0, Real-Time Analysis (RTA) v3.3.3, and bcl2fastq2 software v2.20, converting base calls to FASTQ format. The average percentages of bases achieving Q30 for read 1 and read 2 (mean ± SD) were 94.1 ± 0.1% and 92.7 ± 0.5%, respectively.

RNA-seq reads were mapped to the Canis lupus familiaris reference genome (NCBI Taxonomy ID: 9615, CanFam3.1 assembly, April 2018) using STAR v2.5.2b and Genedata Profiler Genome v10.1.15a. The mean ± SD total reads, mapped reads, and mapping percentages were 59,413,484.8 ± 11,379,311.2, 57,909,272.9 ± 11,119,490.9, and 97.5 ± 0.1%, respectively.

### Effect of NB-UVR on cell viability and apoptosis in canine mast cell line

#### Effect of NB-UVR on Cell Viability and Apoptosis in Canine Mast Cell Line

The canine cutaneous mast cell tumor cell line HRMC [18] was used for in vitro experiments. Cells were cultured in RPMI 1640 medium (Wako Pure Chemical Industries, Osaka, Japan) supplemented with 10% fetal bovine serum (Biowest, France) and 1% penicillin/streptomycin (Sigma Aldrich). Cultures were maintained at 37 °C in a humidified atmosphere containing 5% CO₂.

#### Cell Viability Assay

The effect of NB-UVR irradiation on HRMC cell viability was assessed using the WST assay with a Cell Counting Kit-8 (Dojindo Laboratories, Kumamoto, Japan). Cells were seeded at a density of 1×10^6^ cells/mL in 96-well plates and irradiated with NB-UVR at various doses (100–600 mJ/cm²) and wavelengths (310, 320, and 330 nm). After 24 hours of incubation, CCK-8 reagent was added to each well, and plates were incubated for an additional 4 hours. Absorbance at 450 nm was measured with a microplate reader (Power Scan HT, DS Pharma Biomedical, Tokyo, Japan). Cell viability (%) was calculated using the formula:

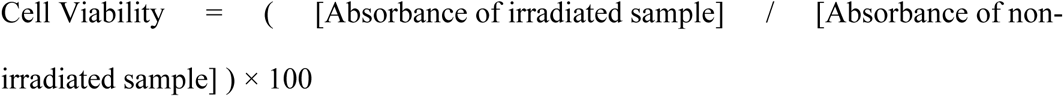

#### Apoptosis Assay

Apoptosis was evaluated using DAPI staining, following a previously described protocol [19]. HRMC cells (1×10^6^ cells/mL) were irradiated with NB-UVR (310, 320, or 330 nm) at 300 mJ/cm². After 24 hours, cells were adhered to slide glasses using LBC Prep 2 (Muto Pure Chemicals), fixed with 4% paraformaldehyde, and permeabilized with 0.2% Triton X-100 (Kanto Chemical, Tokyo, Japan). DAPI solution (VECTASHIELD Mounting Medium with DAPI, Vector Laboratories, Burlingame, CA, USA) was applied to the slides. Apoptotic cells were identified based on nuclear fragmentation and chromatin condensation using a fluorescence microscope (AX80N-05, Olympus, Tokyo, Japan). Five fields per treatment condition were analyzed, and experiments were performed in triplicate.

### Statistical analysis

Statistical analyses were performed using StatMate V software (ATMS, Tokyo, Japan). One-way repeated measures ANOVA was used to evaluate clinical scores, IDT scores, MC numbers in each treatment. One-way ANOVA was used to analyze in vitro experiments of cell viability and the rate of apoptosis in each treatment. Significant differences were further analyzed using Dunnett’s test to compare NB-UVR-irradiated groups with non-irradiated controls. Comparisons among NB-UVR wavelengths (310, 320, and 330 nm) were conducted using Tukey’s multiple comparison test A p-value of <0.05 was considered statistically significant.

Low read counts (<100 total counts per gene) were excluded from RNA-seq data. Normalization and differential expression analysis were performed using the TCC baySeq R package [20, 21], which applies Bayesian statistical methods to minimize false positives and rank differentially expressed genes (DEGs). DEGs were identified at a false discovery rate (FDR) threshold of q < 0.1. Pairwise comparisons were visualized using distance matrices based on Spearman rank correlation in Heatmapper (http://www.heatmapper.ca/pairwise/). Gene Ontology (GO) enrichment (Biological Process Level 3) and KEGG pathway analyses were conducted using DAVID Bioinformatics Resources 2021 (https://david.ncifcrf.gov/) with an FDR threshold of q < 0.1. RNA-seq data have been submitted to the DDBJ; the accession number is forthcoming.

## Results

### Clinical trials in dogs with spontaneous AD

By the fourth treatment session, significant differences in lesional scores were observed among the groups. The 320 nm irradiation group showed a significant improvement compared to the untreated control group (n = 6, p < 0.05; Figs 1A and 1B). No clinical adverse reactions were observed in any of the dogs throughout the study period.

**Fig 1.**
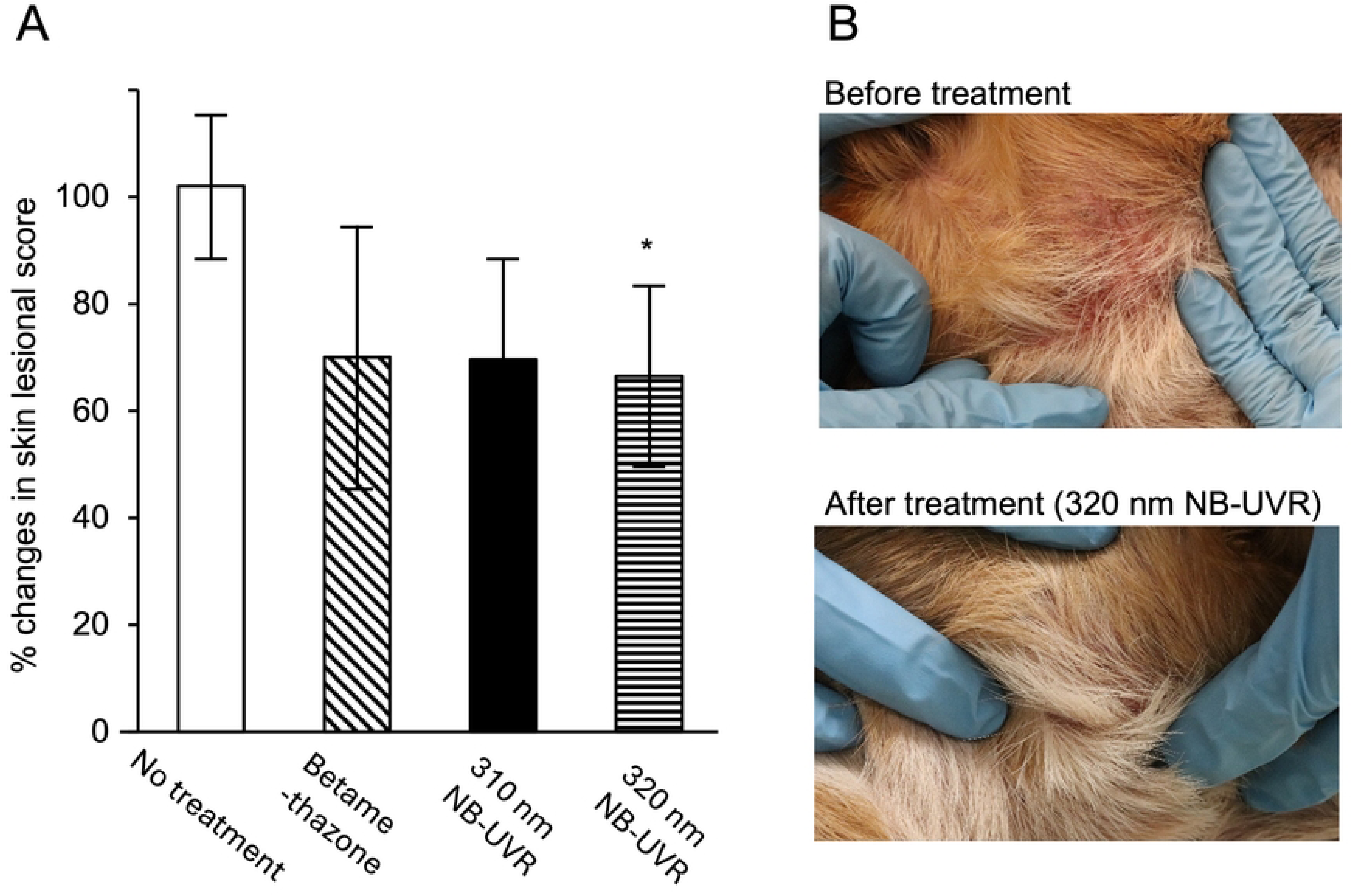
Effect of NB-UVRs on skin lesions in canine atopic dermatitis. (A) Changes in skin lesional scores after the fourth treatment with NB-UVRs or topical ointment. The NB-UVR dose was adjusted to 600 mJ/cm². *Indicates a significant difference compared to the untreated control (n = 6, p < 0.05). (B) Representative clinical findings of a dog treated with 320 nm NB-UVR, showing hair regrowth and reduced erythema in the cervical skin four weeks after treatment.

### Intradermal skin test after NB-UVR irradiation in Beagles

Significant differences in IDT scores were observed for reactions induced by HCO-60 (1×10^6^ v/v) and histamine across all NB-UVR wavelengths compared to the unirradiated control (n = 3, p < 0.01). Furthermore, at 320 nm, a significant decrease (p < 0.05) in IDT score wasin response to the 1×10^5^ v/v of HCO60. (Fig 2).

**Fig 2.**
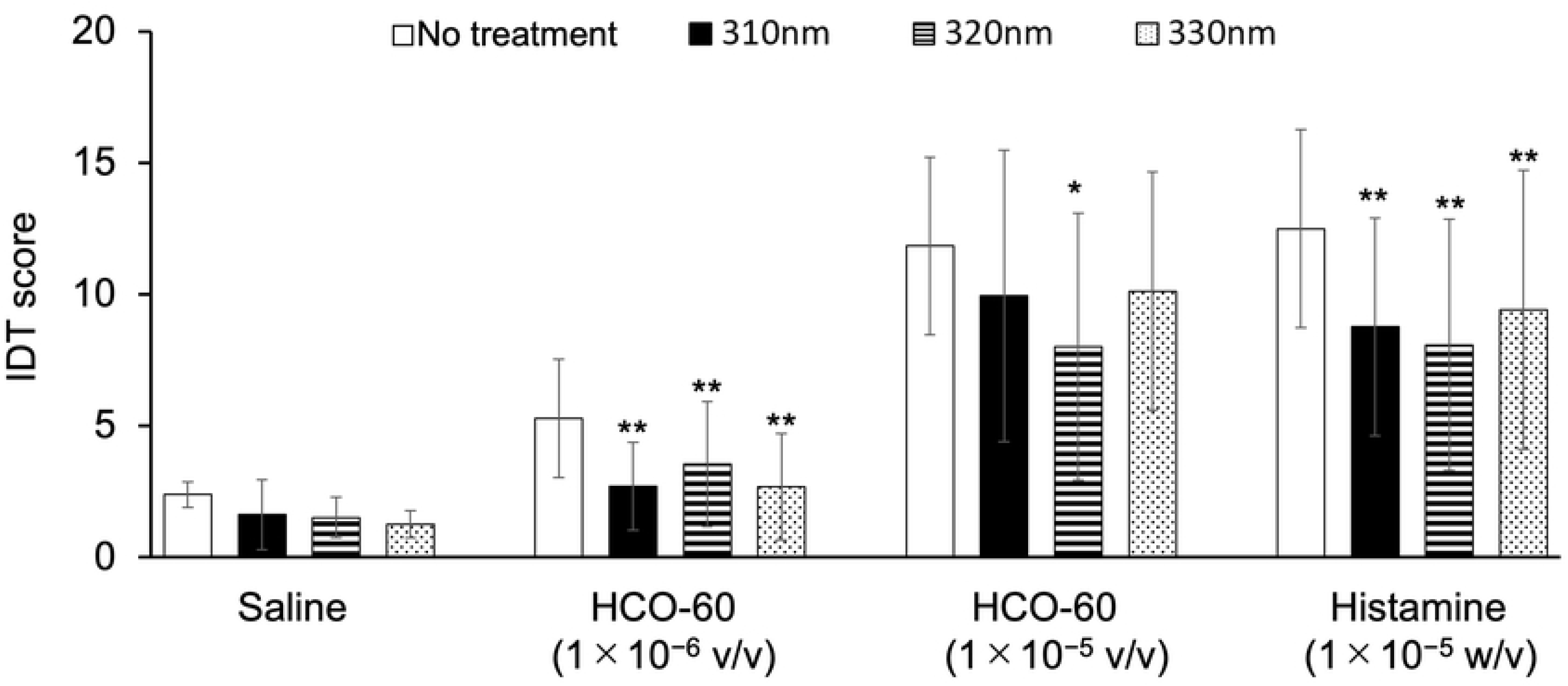
Effect of NB-UVRs on polyoxyethylene hydrogenated castor oil 60 (HCO-60)- and histamine-induced wheal-and-flare reactions in dogs. NB-UVR was applied at a dose of 300 mJ/cm². **Indicates a significant difference compared to the non-irradiated control *(p < 0.05) and **(p < 0.01), n = 3. Data are presented as mean ± standard deviation.

### Histopathological analysis following NB-UVR irradiation in Beagles

Sunburn cells, which represent keratinocytes undergoing apoptosis due to physiological UVR damage [22], were not detected in any of the groups receiving 300 mJ/cm² NB-UVR irradiation. However, irradiation at 310 nm with a higher dose (1500 mJ/cm²) induced visible sunburn cells (S2 Fig). Mast cell (MC) counts in skin sections revealed a significant reduction in MC numbers in the 310 nm NB-UVR groups compared to the control group (p <0.05; Fig 3).

**Fig 3.**
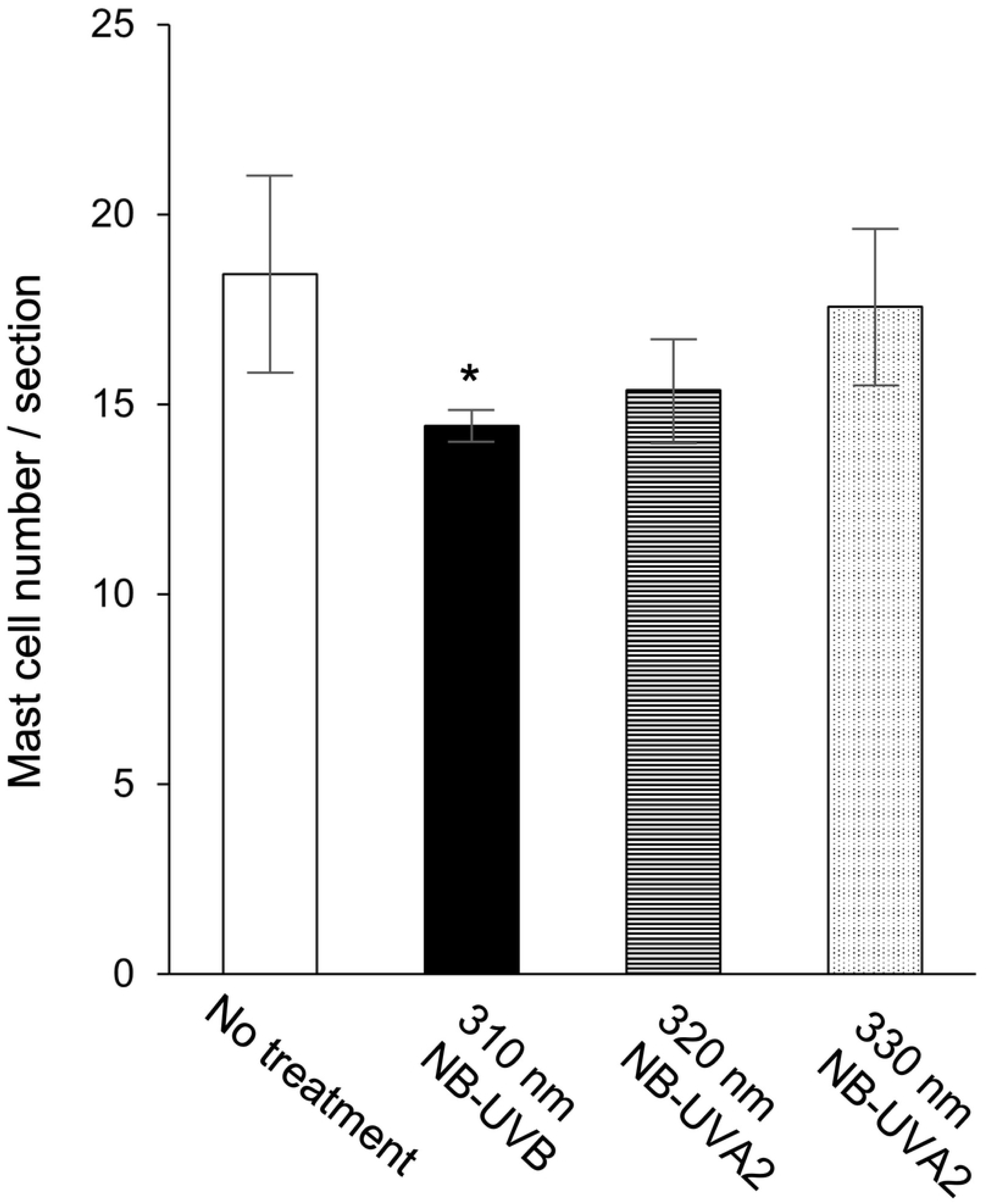
Effect of NB-UVRs on skin mast cell numbers. NB-UVR was applied at a dose of 300 mJ/cm². *Indicates a significant difference compared to the non-irradiated control (n = 3, p < 0.05). Data are presented as mean ± standard deviation.

### Cell viability and apoptosis in HRMC cells

The effects of NB-UVR irradiation on cell viability and apoptosis were evaluated using HRMC cells. Irradiation at 310 and 320 nm NB-UVR dose-dependently reduced HRMC cell viability, while no significant effects were observed with 330 nm NB-UVR (Fig 4A). Within the dose range of 200–600 mJ/cm², 310 nm NB-UVR significantly reduced cell viability compared to 320 and 330 nm. Additionally, cells treated with 310 nm NB-UVR exhibited a significantly increased rate of apoptosis (p < 0.01; Fig 4B).

**Fig 4.**
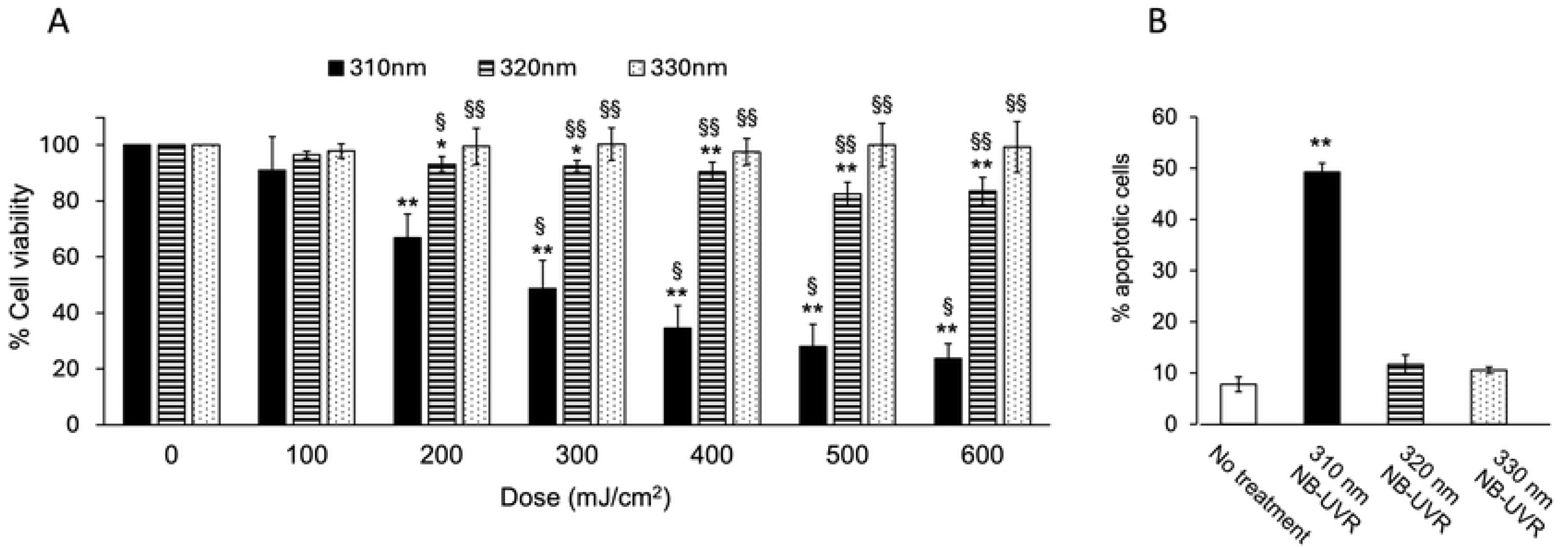
Effects of NB-UVRs on cell viability and apoptosis in HRMC cells. (A) Cell viability assessed using the WST assay. (B) Rate of apoptotic cells determined by DAPI staining. The NB-UVR dose was adjusted to 300 mJ/cm² for apoptosis assessment. *(p < 0.05) and **(p < 0.01) Indicates a significant difference compared to non-irradiated cells. §(p < 0.01) and §§(p < 0.001) Indicates a significant difference compared to cells treated with 310 nm or 320 nm of NB-UVR. Data are presented as mean ± standard deviation.

### RNA-seq analysis after NB-UVR irradiation in Beagles

Out of a total of 15,399 expressed genes, after filtering out 39 non-differentially expressed genes (DEGs), 88 statistically significant DEGs (q < 0.1) were identified using TCC-baySeq analysis (S4 Table). Of the 88 DEGs, the numbers for each NB-UVR irradiated group comparison (300 mJ/cm^2^, 320 nm > others; 1500 mJ/cm^2^, 310 nm > others; others > 1500 mJ/cm^2^, 310 nm; others > 300 mJ/cm^2^, 310 nm; and others > un-irradiated control) were 71, 13, 2, 1, and 1, respectively. Pairwise comparison revealed two distinct clusters separating groups receiving 300 mJ/cm^2^, 320 nm > others and 1500 mJ/cm^2^, 310 nm > others (Fig 5).

**Fig 5.**
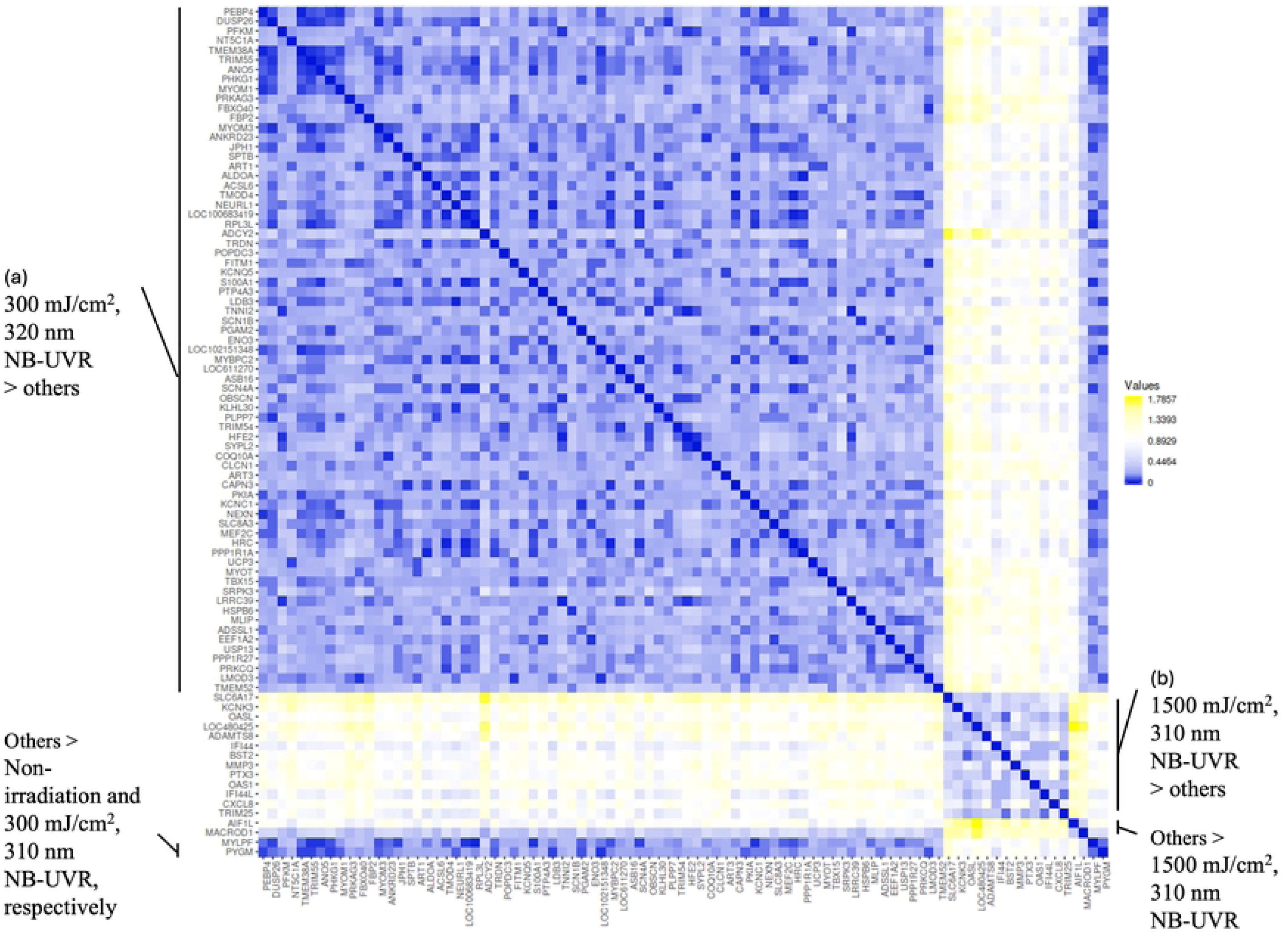
Pairwise comparison of DEGs identified after NB-UVR irradiation (a) DEGs identified following 300 mJ/cm², 320 nm NB-UVA2 irradiation. (b) DEGs identified following 1500 mJ/cm², 310 nm NB-UVB irradiation. These clusters represent the most differentially expressed genes compared to other conditions, as determined by TCC baySeq analysis (FDR < 0.1).

A summary of functional enrichment analysis for DEGs (q < 0.1) using DAVID is presented in Table 1. Irradiation with 300 mJ/cm^2^, 320 NB-UVR enriched biological processes (BP) including **muscle contraction** (GO:0006936), **canonical glycolysis** (GO:0061621), and **fructose 1,6-bisphosphate metabolic process** (GO:0030388). Cellular components (CC) enriched included **M-band** (GO:0031430), **sarcoplasmic reticulum** (GO:0016529), **Z-disc** (GO:0030018), and **axon** (GO:0030424). KEGG pathway analysis indicated involvement in **cytoskeletal regulation in muscle cells** (cfa04820), **glucagon signaling pathway** (cfa04922), **glycolysis/gluconeogenesis** (cfa00010), **adrenergic signaling in cardiomyocytes** (cfa04261), **biosynthesis of amino acids** (cfa01230), **pentose phosphate pathway** (cfa00030), and **fructose and mannose metabolism** (cfa00051).

**Table 1.**
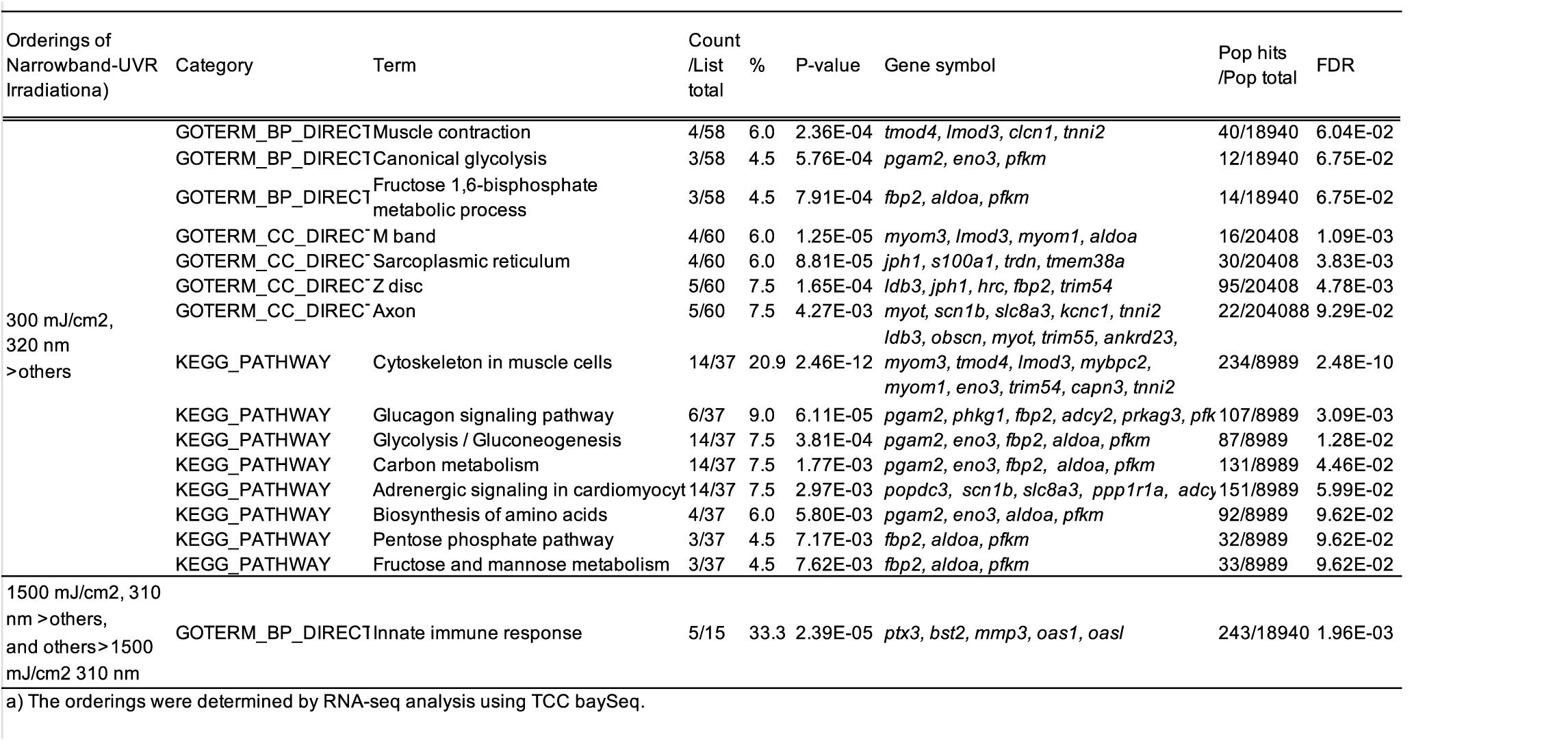
Results of the enrichment analysis (David) of differentially expressed genes in the whole RNA transcriptome of narrowband-UVR irradiated skin analyzed by TCC baySeq.

In contrast, irradiation with 1500 mJ/cm^2^, 310 nm NB-UVR enriched only a single BP term, **innate immune response** (GO:0045087).

RNA-seq data are detailed in S4 Table and have been submitted to the DDBJ; accession numbers will be provided upon publication.

## Discussion

This study explored the non-inflammatory mechanisms of narrowband ultraviolet radiation (NB-UVR) therapy by applying NB-UVR LED devices to a canine model of spontaneous atopic dermatitis (AD) and healthy Beagles. The results highlighted the therapeutic potential of 320 nm NB-UVR irradiation in canine AD and demonstrated the effects of sub-MED NB-UVR on mast cells (MCs) and gene expression. Specifically, NB-UVR suppressed intradermal skin test (IDT) reactions, and the 310 nm UVR reduced MC counts in skin tissue. In vitro studies further revealed that NB-UVR (310 and 320 nm) decreased MC viability, with 310 nm inducing apoptosis.

While guidelines exist for treating spontaneous AD in dogs [23], reports on NB-UVR therapy remain limited. Previous studies using 311 nm NB-UVR in 32 dogs reported MED values of ≥ 432 mJ/cm² [24], while 308 nm NB-UVR in 7 Beagles showed MED values of ≥560 mJ/cm² [25]. Based on our findings (S3 Table) and previous studies, we determined 300 mJ/cm² of 310 nm NB-UVR as the sub-MED dose. Previous research demonstrated that 308 nm NB-UVR irradiation in DNCB-induced dermatitis induces apoptosis and reduces INF-γ and TNF-α levels in skin cells [25], suggesting that UVR effects can penetrate into the dermis, consistent with our in vivo results. Additionally, our in vitro study showed that 310 nm NB-UVR irradiation induced apoptosis in HRMC cells, aligning with findings from previous studies on human mast cells [26]. In our clinical study, the therapeutic efficacy of NB-UVR for canine AD was observed only with 320 nm irradiation. The difference between the effects of 310 and 320 nm may be attributed to erythema induced by 310 nm NB-UVB (600 mJ/cm²), which could influence its therapeutic efficacy compared to 320 nm.

Previous studies have reported an increase in cutaneous mast cell (MC) numbers and histamine release following UVR exposure [5–7, 10], which contrasts with our findings. The UVR doses used in those studies were at least one MED or higher, whereas our study focused on sub-MED doses (300 mJ/cm²). These prior studies highlighted the inflammatory effects of UVR exposure, emphasizing the role of higher doses in inducing such responses.

Patra et al. [27] examined the effect of skin microbiome on UVR-induced immunosuppression against CHS using a single 618 mJ/cm² UVR exposure, which exceeded the minimum inflammatory dose (316 mJ/cm²). Their microarray analysis of whole-transcriptome RNA identified the differentially expressed genes (DEGs) presence or absence of skin microbiome, one of which was the upstream regulator gene IL-1β.

These findings align with our results at 310 nm NB-UVR and 1500 mJ/cm² irradiation (Table 1), where we observed DEGs associated with type-I interferon (IFN1) pathways. Specifically, TRIM25, which interacts with the IFN1-inducible gene RIG-1 [28], was upregulated. RIG-1, a cytosolic pattern recognition receptor (PRR), recognizes short double-stranded RNA (dsRNA) generated after UVB exposure [29]. Other IFN1-related genes, such as OAS1, BST2, and IFI44, were also identified [30]. CXCL8/IL-8, a key component of the RIG-I-like receptor signaling pathway (cfa04622), and MMP3, a UVR-responsive matrix metalloproteinase, were similarly upregulated [31]. Of particular interest, OAS1 requires dsRNA as an obligatory cofactor for interferon induction [32], suggesting that dsRNA could function as one of the chromophores in our study. These results underscore the complex interplay between UVR exposure, gene expression, and the immune response, particularly in the context of sub-MED and high-dose UVR irradiation.

Few studies have investigated the effects of sub-MED UVR irradiation. Danno et al. [8] demonstrated that UVR exposure at 50–100 mJ/cm², with a peak emission at 305 nm (wavelength range: 280–370 nm), suppressed IDT reactions and mast cell (MC) degranulation in mice. Similarly, Byne et al. [9] reported dose-dependent suppression of CHS with UVR irradiation (260–320 nm) at doses below the minimum edematous dose (35–280 mJ/cm²), administered daily for three days. Our findings with 310 nm UVR at 300 mJ/cm² were consistent with these studies, while providing greater specificity due to the narrower half-bandwidth of 10 nm.

Byne et al. [9], also reported different dose-response effects for UVA (320–400 nm) compared to UVB, which aligns with our observations at 320 nm NB-UVR. The MED of 320 nm UVR in our study (>1500 mJ/cm²) closely matched the minimal edematous dose of UVA reported by Byne et al. (3360 mJ/cm²). Furthermore, Danno et al. [8] suggested that UVB exerts dual effects on MCs, depending on the dose. UVB doses exceeding 200 mJ/cm² induced inflammatory responses, while sub-MED UVB altered the MC/vasoactive amine system, suppressing ear swelling in response to degranulators. Similarly, our results suggest that 320 nm NB-UVR irradiation elicited a non-inflammatory response to UVR exposure.

To our knowledge, no previous studies have systematically compared irradiation effects using such carefully controlled half-bandwidths within a 10-nm range. Further research is warranted to explore the biological effects of UVR doses in the border region between UVA and UVB, particularly across both low- and high-dose ranges.

RNA-seq analysis revealed that 320 nm NB-UVR irradiation upregulated genes associated with muscle function. The arrector pili muscle (APM), a smooth muscle located in the upper and middle dermis, connects the epidermal basement membrane to the bulge region of hair follicles (S2 Fig) and may be influenced by UVR exposure. While common APM markers such as ***Acta2***, ***Cnn1***, ***Myh11***, and ***Tagln*** were not detected among the differentially expressed genes (DEGs) following 320 nm irradiation, other markers, ***Hspb6*** and ***Nexn***, identified in single-cell RNA-seq studies, were upregulated [33–35]. ***Hspb6*** encodes HSP20, a heat shock protein highly expressed in smooth muscle cells. Beta-adrenergic agonists stimulate HSP20 phosphorylation via cAMP, which decreases intracellular calcium levels and promotes the dephosphorylation of 20 kDa myosin light chains, ultimately inhibiting smooth muscle contraction [36]. ***Nexn*** encodes Nexilin, a protein localized to dense bodies and bands in smooth muscle, playing a key role in actin polymerization, cell migration, and differentiation, as demonstrated through gene silencing experiments in smooth muscle cells [37].

The DEGs (*Popdc3*, *Scn1b*, *Slc8a3*, *Ppp1r1a*, *Adcy2*) identified in the 320 nm irradiation group were associated with the adrenergic signaling in cardiomyocytes pathway (cfa04261). These genes encode proteins such as adenylate cyclase (AC), ion channels, and a serine/threonine phosphatase inhibitor, suggesting involvement in G-protein coupled receptor (GPCR) signaling.

Recent research by Schwartz et al. (2020) revealed that sympathetic nerves innervate both the arrector pili muscles (APMs) and hair follicle stem cells (HFSCs)

[38]. Piloerection, triggered by APM contraction, maintains sympathetic innervation to HFSCs. The **β2-adrenergic receptor (ADRB2)** in HFSCs directly responds to norepinephrine and cold signals from the nervous system, activating muscle contraction and initiating a new hair cycle via the GPCR/Gαs/AC/cAMP/CREB pathway [39]. RNA-seq experiments in mice, comparing genes expressed after treatment with either cAMP or forskolin, and those in ***Adrb2***-conditionally deleted HFSCs, identified many genes involved in glycolytic metabolism that were upregulated by CREB activation [39], These include *Fbp2*, *Aldoa*, *Pfkm*, *Pgam2*, and *Eno3*, which were also enriched in the KEGG pathway associated with 320 nm irradiation in our study (Table 1). This suggests that 320 nm irradiation may induce HFSC changes similar to those triggered by cold stimulation.

The detection of axon-related components (GO:0030424) in our data further supports this hypothesis (Table 1). Cold exposure-induced piloerection typically enhances sympathetic nerve activity, resulting in increased norepinephrine release, elevated heart rate, and glucagon secretion. Elevated glucagon levels, in turn, stimulate gluconeogenesis in the liver, producing heat [40]. The KEGG pathways identified in our study may reflect similar systemic adrenergic responses induced in the skin by 320 nm irradiation.

The transcription factor-encoding genes ***Mef2c*** and ***Tbx15*** were among the differentially expressed genes (DEGs) upregulated by 320 nm irradiation (S4 Table). **Myocyte Enhancer Factor 2C (MEF2C)** is a critical regulator of muscle and nervous system development and function [41, 42]. MEF2C directly upregulates the expression of ***Hrc***, which encodes histidine-rich calcium-binding protein (HRCBP). HRCBP is localized to calciosomes within arterial smooth muscle cells [43] and interacts with triadin in the sarcoplasmic reticulum [44]. Both HRCBP and triadin play essential roles in calcium signaling pathways (cfa04020), influencing muscle contraction by regulating intracellular calcium concentrations.

**T-box transcription factor 15 (TBX15)** affects hair length, pigmentation, and the musculoskeletal system in mice [45]. In mouse skin at postnatal day 3.5, ***Tbx15*** is strongly expressed in the condensed upper dermis and the developing dermal sheaths of hair follicles, with lower levels detected in both dorsal and ventral skin.

The upregulation of ***Mef2c****, **Hrc**, **Trdn**,* and ***Tbx15*** in our study suggests that 320 nm irradiation promotes cellular growth and differentiation processes, particularly those related to muscle function and hair follicle development.

Our findings demonstrated dual effects of NB-UVB irradiation: inflammatory responses at 310 nm (300 and 1500 mJ/cm²) and anti-allergic, anti-IDT effects, along with gene expression changes associated with arrector pili muscle (APM) function and hair follicle stem cell (HFSC) regeneration at 320 nm (300 mJ/cm²). These results uncover a novel mechanism of NB-UVB phototherapy, emphasizing the importance of dose- and wavelength-dependent effects.

## Acknowledgement

The authors extend their sincere gratitude to Dr. Keitaro Ohmori from the Cooperative Division of Veterinary Sciences, Graduate School of Agriculture, Tokyo University of Agriculture and Technology, for generously providing the HRMC cell line. We would like to thank Dr. Azusa Ogita, and Dr. Keigo Ito (Division of Dermatology and Dermatopathology, Nippon Medical School, Musashi Kosugi Hospital), and Dr. Hidenori Matsuda (Shibuya Aesthetic Surgery Clinic), taught me the knowledge and skills of dermatology practice, and Hitomi Endo for her assistance of animal experiments.

## Supporting information

S1 Fig. Spectra of narrow band UVR devices

S2 Fig. Histopathological findings of NB-UVRs irradiated canine skin. Inflammatory cell infiltrate to the superficial dermis and apoptotic keratinocytes (arrow) are observed in 1500 mJ/cm^2^, 310 nm NV-UVR irradiated canine skin. (HE staining x100)

S1 Table. Profiles of dogs with spontaneous atopic dermatitis used in the NB-UVRs clinical trial

S2 Table. A dataset of skin lesions treated in each case

S3 Table. Minimal erythema dose in six healthy beagles

S4 Table. Diffrentially expressed gene of NB-UVRs shown by RNA-seq analysis using TCC baySeq

